# Whose sample is it anyway? Widespread misannotation of samples in transcriptomics studies

**DOI:** 10.1101/064626

**Authors:** Lilah Toker, Min Feng, Paul Pavlidis

## Abstract

Concern about the reproducibility and reliability of biomedical research has been rising. An understudied issue is the prevalence of sample mislabeling, one impact of which would be invalid comparisons. We studied this issue in a corpus of human genomics studies by comparing the provided annotations of sex to the expression levels of sex-specific genes. We identified apparent mislabeled samples in 46% of the datasets studied, yielding a 99% confidence lower-bound estimate for all studies of 33%. In a separate analysis of a set of datasets concerning a single cohort of subjects, 2/4 had mislabelled samples, indicating laboratory mix-ups rather than data recording errors. While the number of mixed-up samples per study was generally small, because our method can only identify a subset of potential mix-ups, our estimate is conservative for the breadth of the problem. Our findings emphasize the need for more stringent sample tracking, and that re-users of published data must be alert to the possibility of annotation and labelling errors.

---

Recent years have seen an increase in concern about the quality of scientific research, along with efforts to improve reliability and reproducibility [1,2]. These issues are highly relevant to genomics studies, which deal with complex and often weak signals measured genome-wide. In transcriptomics studies (our focus here), mRNA is extracted from samples and processed using microarrays or RNA-seq, followed by statistical analysis to identify patterns of interest (e.g. differential expression). Much work has been done to raise awareness of technical issues in such studies such as RNA quality [3] and batch effects [4] and many investigators are aware of the need to address them [5]. In addition, great effort was put into establishing guidelines for annotation standards of expression data into public repositories [6] In contrast, a key step in many scientific experiments, but which has received less attention, is the importance of maintaining an accurate correspondence between the experimental conditions or sources of the samples and the eventual data. Simply put, for the analysis to be valid, the samples must not be mixed up. If mix-ups are present but undetected, the conclusions of the analysis might be affected and pollute the literature, as well as create a lurking problem for those who re-use the data.

The obviousness of the need to avoid mix-ups suggests that investigators should be well aware of the risk, and take steps to reduce it, such as careful bookkeeping (e.g., permanent sample tube labels matched to data files). However we recently became concerned that mix-ups might not be rare. Our concerns came to a head when we reanalyzed four publically available datasets of Parkinson’s disease subjects [7]. As part of our quality checks of the data, we examined expression levels of sex-specific genes (genes expressed only in males or in females), and compared these with the corresponding subject sex meta-data annotations from each of the papers. To our surprise, we found discordance between the sex predicted based on expression levels of sex-specific genes and the manuscript-annotated sex in two out of the four datasets (Supplementary Fig. S3). This finding, and other anecdotal observations, led us to examine this issue more broadly.

Sex-specific genes are well-suited for this purpose. In genetics studies, genotypes of the sex chromosome are routinely used to identify mislabeled samples [8,9], moreover, sex check is a built-in option for some of the dedicated software [10]. Given that genetic abnormalities resulting in disagreement between genotypic and phenotypic sex are rare [11], any disagreements are very likely to stem from errors and may also be indicative of other dataset quality issues. Using such genes for quality checks of transcriptome data is not widespread practice, but it is well known that several X- and Y-linked genes show sex-specific patterns of expression. A limitation of this approach is that mix-ups that do not yield conflicting sex labels (e.g., swapping two female samples) cannot be detected. But at the very least it can provide a lower bound for the amount of mix-ups and if any are detected it should trigger a reassessment of the tracking of all samples in the study.

In this study, we focused on publically available human expression profiling experiments that included individuals of both sexes. To our surprise we found strong evidence of mix-ups in nearly half of them. Importantly, for the vast majority of the studies we were able to validate that the disagreement between metadata- and gene-based gender is prevalent in the original manuscript. This indicate that the disagreements are not a result of erroneous gender description during data submission to public repository. An additional 10% of the studies have samples of ambiguous sex that suggests the possibility of samples being mistakenly combined or other quality problems. While it is possible that a small number of the cases we identify are due to sex chromosome abnormalities, we regard the most likely explanation for most to be laboratory mix-ups or errors in the meta-data annotations. Our findings suggest a widespread quality control issue in genomics studies.

## Results

We identified a corpus of 70 human gene expression studies that had sample sex annotation (4160 samples in total) run on two platforms. We developed a simple robust method for classifying samples by sex based on three sex specific genes – *XIST*, *RPS4Y1* and *KDM5D*. *XIST* (*X-inactive specific transcript*) is expressed from the inactive X chromosome and acts to silence its expression and thus, is only expressed in female subjects. *KDM5D* (*Lysine (K)-Specific Demethylase 5D*) and *RPS4Y1* (*Ribosomal Protein S4, Y-Linked 1*) are both located on the Y chromosome, and thus are only expressed in male subjects. Although additional sex-specific genes exist, we determined that *KDM5D, RPS4Y1* and *XIST* are the only sex-specific genes consistently showing high expression levels in the associated sex in all tissues. Our method assigns a predicted sex based on gene expression to each sample, which we refer to as “gene-based sex” (see methods). We also performed a second analysis to identify samples where a gene-based sex could not be confidently assigned. Such samples might reflect technical problems, but could also be due to true biological effects; for example, *XIST* expression is altered in some cancers and in early stages of development [12]. We then compared gene-based sex to the sex according to the provided sample annotations (“meta-data-based sex”) for the 70 studies, seeking samples with disagreements. Fig. 1 shows examples of studies with no discrepant samples (1A) and with discrepancies (1B). Similar plots for all datasets analyzed are shown in supplementary Fig. S2. All calls of discrepant or ambiguous sex were followed by manual confirmation.

**Figure 1.**
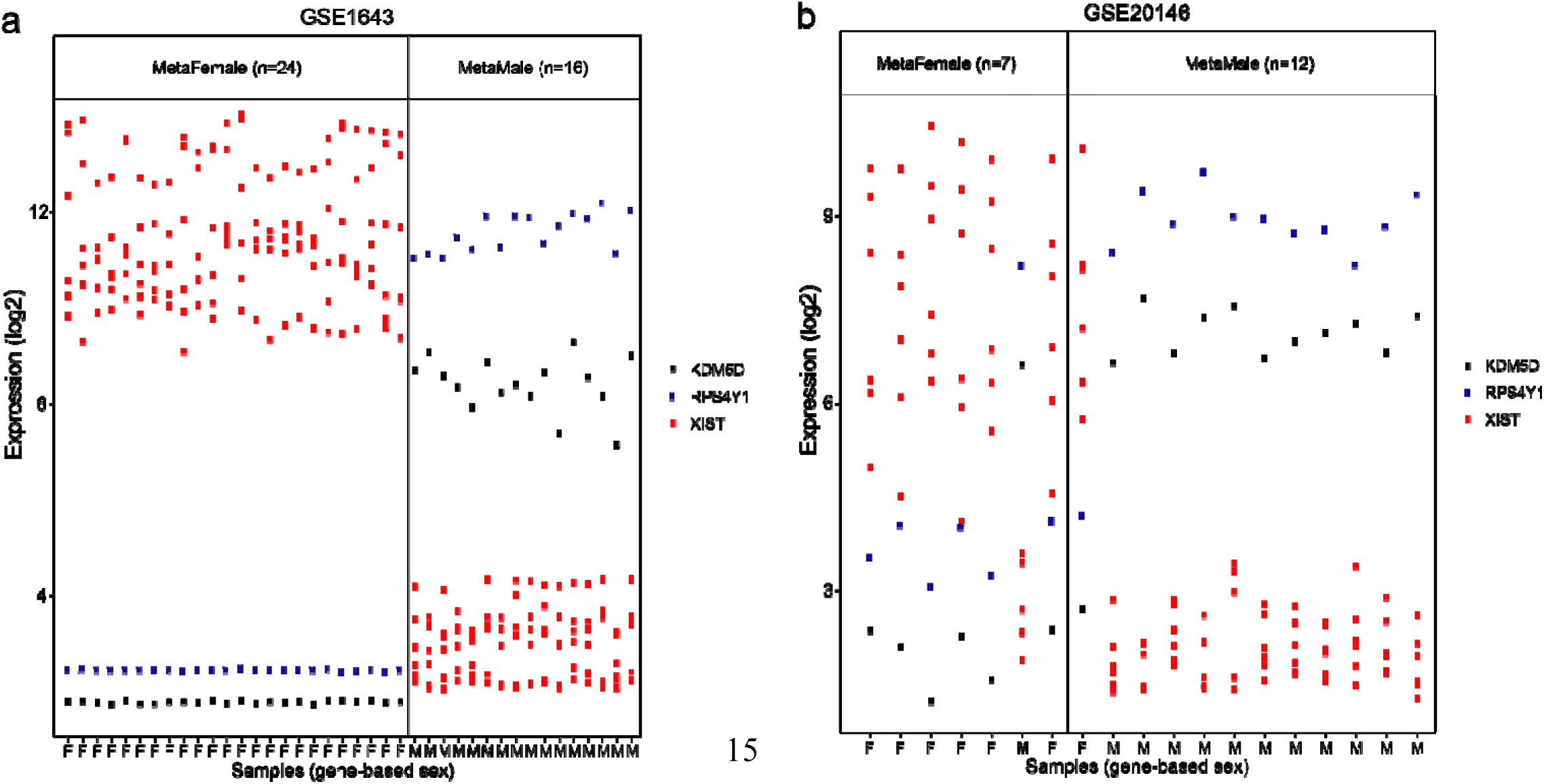
Representative plots showing expression levels of sex-specific probesets. Expression level of probesets representing the *XIST*(red), *KDM5D* (black) and *RPS4Y1* (blue) genes. “MetaFemale” and “MetaMale” indicate the meta-data annotated sex of the samples and their total number in brackets. The “M” and “F” along the X axis indicates the gene-based sex of the samples, as determined by k-means clustering. Log_2_-transformed expression levels are plotted. **(a)** Representative dataset with no mismatched samples. **(b)** Representative dataset with two mismatched samples (highlighted with grey bars). Gene-based sex that contradicts the annotated sex of the sample is highlighted in bold at bottom.

We found samples with a discrepancy between the meta-data sex information and the gene-based sex in 32/70 (46%) of the datasets (ambiguous samples excluded; summarized in Table 1; details in supplementary table S2). Although datasets containing mismatched samples were more prevalent among cancer datasets (50% vs 43%, cancer vs. non-cancer, respectively), the proportion of mismatched samples was similar in cancer and non-cancer samples (2.14% vs 1.93%; Table 1). This discrepancy might be explained by on average higher number of samples in cancer datasets from our corpus (Supplementary Table S1). As expected, the proportion of samples with ambiguous gene-based sex was much higher in cancer as compared to non-cancer samples: 23/887 (2.6%) in cancer vs. 11/3273 (0.3%; Table 1). In total, 34 samples were flagged as ambiguous, though we note that 12/34 (35%) would have been signed to the discrepant sex by our method. Ambiguous samples were found in 15/70 (21%) of the studies (eight of which also contained mismatched samples).

**Table 1.** Summary of discrepancies between the gene expression-based and annotated sex in human microarray datasets.

|  | All Datasets | Non-cancer Datasets | Cancer Datasets | All Samples | Non-cancer Samples | Cancer Samples |
| --- | --- | --- | --- | --- | --- | --- |
| Correctly annotated | 31 (44%) | 30 (51%) | 1 (8%) | 4043 (97%) | 3213 (98%) | 830 (94%) |
| Mismatched | 31 (46%) | 25 (43%) | 6 (50%) | 83 (2%) | 64 (1.9%) | 19 (2.14%) |
| Unclassified | 15 (21%) | 7 (12%) | 8 (67%) | 34 (0.8%) | 11 (0.3%) | 23 (2.6%) |
| Total | 70 | 58 | 12 | 4160 | 3273 | 887 |

Unclassified samples are samples with disagreement between their classification using k-means clustering and the median expression of the sex specific probesets.
Datasets were considered as “correctly annotated” only if they did not contain mismatched or unclassified samples

Because the sex annotations we used to this point were obtained from the sample descriptions in GEO, there was a possibility that the discrepancies we identified were due to mistakes introduced during the communication of the data from the submitter to GEO. If this were the case, the results in the original publication (29/31 of the affected studies had an associated publication) would be unaffected, though users of the GEO data would still be affected. To check this possibility, we went back to the 29 original publications to see if the sex labels provided in the paper matched those in GEO (detailed in Supplementary Table 1). This check was not always possible because many publications did not provide detailed meta-data in the paper or supplementary materials; GEO provides the only record. In 12/31 cases, sufficient detail was provided for us to confirm that the discrepant sex labels were present in the publication, and in all of them there was agreement between the meta-data in the publication and the meta-data in GEO. In 13 cases only summaries were given in the publication (e.g. “10 males and 9 females in group X”). In 10 of these 13 studies, the summary counts in the publication agree with GEO. In the other three, both GEO and gene-based totals disagree with the publication-based totals. In other words, there seems to have been miscommunication with GEO in addition to a sex annotation discrepancy in the original study report. Finally, for four data sets meta-data was not provided or ambiguously described in the paper. We failed to find any unambiguous case in which we would infer the only problem was a miscommunication with GEO.

The analysis presented cannot distinguish between actual sample mix-ups (e.g., tube swaps) and errors in the meta-data (incorrect recording of the subject’s sex). Fortuitously, we identified data sets where it can be determined that at least in some cases, samples were probably physically mixed up. In addition to the 70 data sets used above, we analyzed four datasets that all used human brain RNA from the same collection of subjects (the Stanley Array Collection http://www.stanleyresearch.org/brain-research/array-collection/). In this case the meta-data is common across the four laboratories since they are all analyzing the same individuals (though not all studies analyzed all the individuals). If the meta-data is incorrect, then all of the studies should show discrepancies for the same samples. If the samples were mixed up in a particular laboratory (or by the sample provider at the time they were sent to the laboratory), each study would have different discrepancies. We found that out of the four available datasets with data corresponding to the same subjects, two datasets contained mismatched samples (a single mismatched sample was identified in the “AltarA” study, and five in the “Dobrin” study; supplementary Fig. S3). Importantly, the mismatched subjects differed between the datasets and samples from the same subjects appeared as correctly annotated in the other datasets. This suggests that the mismatched cases are likely to represent mislabeled samples rather than mistakes taking place during the recording of the subjects’ sex.

We were surprised that nearly 50% of studies had at least one labeling error, and were concerned that this might be an overestimate by chance, due to sampling error. To address this we computed confidence intervals for our estimate of the fraction of affected studies, yielding a 95%-confident lower bound of 36% and a 99% lower bound of 33% (upper confidence bounds were 56% and 60% respectively). We also note that our independent observations of 2/4 datasets containing misannotations described in Santiago et al. [13] and in 2/4 of the Stanley data set are in agreement with a relatively high estimate. Thus we project that, with 99% certainty, if all expression studies in GEO could be checked for mix-ups based on sex-specific genes, the fraction affected would be at least 33%.

## Discussion

Using a simple approach to compare sample annotations for sex to expression patterns, we found that nearly 50% of datasets we checked contain at least one discrepancy. Our findings are also in general agreement with another study that examined this issue in cancer datasets [14], although in cancer there is an expectation of more ambiguity of sex marker expression [12]. In the case of the Stanley brain datasets, we could determine that the problem is likely to stem from laboratory mix-ups rather than an error in recording the subject’s sex. While our analysis is limited to a corpus of studies where sex information was available along with the presence of good markers on the microarray platform, our data suggest a widespread problem.

What is the impact of this issue? Viewed optimistically, a single mixed-up sample is not likely to dramatically affect the conclusions of a well-powered study. In addition, our analysis suggests a lower (99% confident) estimate of “only” 33% of studies with a sex misannotation, which might provide a small amount of comfort to optimists – it could be worse. However, the sample mislabeling we identified might be the tip of the iceberg, because sex-specific genes can only reveal mixed-up samples with differing sex. We also suggest that sample mix-ups might correlate with other quality problems. Indeed, many of the misannotated datasets we found have additional issues such as undocumented batch effects, outlier samples, other apparent sample misannotations (not sex-related), and discordance in sample descriptions reported in different parts of the relevant publication (supplementary table S2). The presence of samples with ambiguous gene-based gender in non-cancer samples is suggestive of even more quality problems. This is because expression patterns of sex-specific genes could be treated as a positive quality control for the expression data as a whole, serving as indicators for the reliability of other gene signals. Deviations from the expected pattern might indicate samples were mixed together, or suggest problems with RNA quality.

Our conclusions are two-fold. First, there is an alarming degree of apparent mislabeling of samples in the genomics literature. In at least the specific cases we identified, the trust in the reliability of the findings reported is certainly not improved. Second, because it is simple to check the expression patterns of sex markers, the tests we performed should become a routine part of all genomics studies where sex can be inferred from the data.

## Methods

Except where mentioned, data analysis was performed using the R/Bioconductor environment [15,16]. Source code for the analysis is available in a Github repository (https://github.com/min110/mislabeled.samples.identification).

We identified datasets containing sex information as experimental factors by searching the Gemma database [17]. Out of an initial 121 data sets we focused on 79 studies run on the Affymetrix HG-U1333Plus_2 and HG-U133A platforms as they have the same sex marker genes (GEO platform identifiers GPL570 and GPL96 respectively). The annotations in Gemma, which originate from GEO sample descriptions augmented with manual annotation, were re-checked against GEO, resulting in the correction of errors for 14 samples. Data sets that contained samples of only one sex, represented data from sex-specific tissues (e.g. ovary or testicle) or contained numerous missing values were excluded (9 datasets). A final set of 70 studies (a total of 4160 samples) met criteria. Table 1 summarizes the data included and full details of each study are in Supplementary Table S1. Whenever possible, data were reanalyzed from.CEL files. The signals were summarized using RMA method from the Affymetrix “power tools” (http://media.affymetrix.com/partners_programs/programs/developer/tools/powertools.affx), log_2_ transformed and quantile normalized as part of the general Gemma pre-processing pipeline.

**Probeset selection:** The male-specific genes *KDM5D* and *RPS4Y1* are represented by a single probeset on both platforms included in our analysis. *XIST* is represented by two probesets on the GPL96 platform and by seven probesets on the GPL570 platform. With the exception of the 221728_x_at probeset, *XIST* probesets were highly correlated with each other, and negatively correlated with the *KDM5D* and *RPS4Y1* expression in all of the datasets analyzed (supplementary Fig. S4). The poor-performing *XIST* probeset (221728_x_at) was excluded from further analysis. The final set was 4 probesets for GPL96 and 8 probesets for GPL570.

**Assigning gene-based (biological) sex to samples:** The expression data for the selected sex markers were extracted from the normalized data for each data set. For each of these small expression matrices, we applied standard k-means clustering (using the “kmeans” function from the “stats” package in R [18] to classify the samples into two clusters. We assigned the two clusters as “male” or “female”, based on the centroid values of each of the probesets: specifically, the cluster with higher values of the *XIST* probesets centroids and a lower value of *KDM5D* and *RPS4Y1* centroids was assigned as a “female” cluster. To identify samples with ambiguous sex, we calculated the difference between the median expression level of the *XIST* probesets and the median expression level of the *KDM5D* and *RPS4Y1* probesets. We compared this difference with the cluster-based gender, and validated that the difference is positive for samples assigned as females and negative for samples assigned as males. We excluded 34 samples that showed disagreement in this comparison since they could not provide a conclusive result for the gene—expression-based sex. We note that 12 (35%) of these would have been assigned to a cluster contradicting their annotated sex if we had retained them.

**Manual validation of the discrepancy between the gene-based sex and the meta-data-based sex:** For all the cases where a discrepancy was found between the gene-expression-based sex and the meta-data-based sex, we manually examined the original studies to check if the mismatch was due to incorrect annotation of the sample during the data upload to GEO, or was present in the original paper. Since most of the manuscripts only contain summary statistics of the demographic data (13/31, supplementary table S2), direct sample-by-sample validation was not possible for most studies. For these studies we used the highest resolution level of group summary statistics, provided in the publication to validate that the data in the paper corroborate the data in GEO. In addition, for all of the datasets with mismatched samples, we manually evaluated the expression values of the relevant probesets using the GEO2R tool on the GEO website.

**Confidence interval estimate for population proportion of studies with misannotated samples:** We used the properties of the binomial distribution to compute the confidence interval for the population estimate of affected data sets using the “qbinom” function in R.

**Analysis of Stanley Foundation datasets:** CEL files and sample metadata were downloaded directly from the Stanley Medical Research Institute genomic database (https://www.stanleygenomics.org/stanley/). CEL files were pre-processed, quantile normalized and log_2_ transformed using the rma function from the “affy” package in R Bioconductor [15,16].

## Acknowledgements

We thank Patrick Tan for assistance with Gemma query and Ben Callaghan for assistance with the Stanley data sets.

## Author Contributions

LT and PP conceived the study idea. MF and LT performed the analyses, LT and PP prepared the manuscript.

## Competing financial interests

The authors declare no competing financial interests.

## Supporting information captions

**Table S1. Description of datasets included in the study**

**Table S2. Detailed description of all datasets with mislabeled samples**

**Fig S1. Disagreement between gene-based and annotated sex in three datasets participating in the metaanalysis of Parkinson’s disease**

**Fig S2. Expression of probesets corresponding to the sex-specific genes *XIST, KDM5D* and *RPS4Y1* in datasets analyzed in the current study.**

**Fig S3. Gene-based and metadata-based sex in four datasets of similar subjects from Stanley Array collection.**

**Fig S4. Correlation of probests corresponding to sex-specific genes**

